# ACE2 associates with insulin-responsive GLUT4 dynamics in adipocytes

**DOI:** 10.64898/2026.03.10.710920

**Authors:** Takeru Fukushima, Natsuho Moriyama, Haya Sato, Hafumi Nishi, Gwyn Gould, Makoto Kanzaki

## Abstract

Angiotensin-converting enzyme 2 (ACE2) is expressed in adipocytes, yet the mechanisms regulating its intracellular trafficking remain unclear. Here, we investigated whether ACE2 trafficking is coordinated with insulin-responsive vesicle dynamics mediated by the glucose transporter GLUT4 in 3T3L1 adipocytes. Subcellular localization analyses revealed that adipocyte differentiation promotes partial incorporation of ACE2 into insulin-responsive GLUT4-associated vesicular compartments, whereas ACE2 displayed a diffuse distribution in fibroblasts lacking a mature GLUT4 trafficking system. Reconstitution of insulin-responsive GLUT4 vesicle formation through exogenous expression of Sortilin and AS160 in fibroblasts was sufficient to partially recruit ACE2 into perinuclear GLUT4-positive compartments, indicating dependence on canonical GLUT4 vesicle machinery.

NanoBiT assays demonstrated a regulated association between ACE2 and GLUT4 that was modestly enhanced by acute insulin stimulation but reduced following prolonged insulin exposure. Insulin stimulation also produced a slight increase in ACE2 surface exposure, while association with GLUT4 was accompanied by reduced ACE2 shedding, suggesting that recruitment into distinct trafficking routes may alter ACE2 accessibility to shedding machinery in adipocytes. Structural modeling further suggested that ACE2 and GLUT4 can form a membrane-compatible complex.

Together, these findings indicate that ACE2 trafficking is coordinated with insulin-responsive GLUT4 vesicle dynamics, revealing a previously unrecognized association between metabolic signaling and ACE2 cellular dynamics in adipocytes, with potential implications for metabolic dysfunction and ACE2-associated disease processes.

## Introduction

Angiotensin converting enzyme 2 (ACE2) is highly expressed in adipocytes [1] and plays important roles in metabolic regulation and blood pressure control, acting in part through modulation of the renin–angiotensin system (RAS) [2, 3]. Beyond its metabolic functions, ACE2 has gained considerable attention as the cellular receptor that facilitates SARS-CoV-2 entry and COVID-19 pathogenesis [4]. The strong association between obesity and increased COVID-19 severity [5] underscores the importance of understanding ACE2 biology in adipose tissue [6]. Despite its high expression in adipocytes, however, the mechanisms governing ACE2 localization, trafficking, and membrane organization in these cells remain poorly defined.

GLUT4 is an insulin-responsive glucose transporter predominantly expressed in adipocytes and muscle cells, where it mediates insulin-stimulated glucose uptake [7]. Insulin action critically depends on regulated intracellular trafficking of GLUT4 through specialized GLUT4 storage vesicles (GSVs), whose formation and mobilization involve coordinated vesicular machinery [8]. Emerging evidence suggests a functional connection between ACE2 and insulin sensitivity. ACE2 deficiency exacerbates high-fat-diet (HFD)-induced insulin resistance accompanied by reduced GLUT4 expression [9], whereas ACE2-deficient mice show age-dependent resistance to HFD-induced obesity [10]. These seemingly paradoxical findings highlight the complexity of ACE2 function in metabolic tissues and raise the possibility that ACE2 may intersect with insulin-regulated trafficking pathways that control GLUT4 dynamics.

A precedent for ACE2-dependent regulation of membrane transporters is provided by its interaction with B^0AT1 (SLC6A19), a neutral amino acid transporter of the solute carrier (SLC) family [11, 12]. In intestinal epithelial cells, ACE2 functions as a chaperone required for SLC6A19 surface expression; ACE2 deficiency results in SLC6A19 mislocalization and impaired amino acid absorption [13, 14]. Cryo-electron microscopy studies further demonstrated that ACE2 and SLC6A19 form a stable heterotetrameric complex [15], providing structural insight into ACE2-mediated transporter regulation and its involvement in SARS-CoV-2 binding [16]. Given that adipocytes lack significant SLC6A19 expression, an unresolved question is whether ACE2 interacts with alternative SLC transporters in adipose cells. GLUT4 (SLC2A4), as a highly regulated and insulin-responsive transporter, represents a compelling candidate.

To address this question, we investigated whether ACE2 associates with GLUT4 and becomes incorporated into insulin-responsive vesicular compartments. We then performed a series of cell-biological analyses in 3T3-L1 adipocytes to examine ACE2 subcellular localization and insulin responsiveness. We evaluated proximity-based association between ACE2 and GLUT4 using the split luciferase (NanoBiT) system [17] and conducted structural modeling with AlphaFold2 [18]. Our findings suggest that ACE2 forms an insulin-regulated complex with GLUT4, is partially recruited into GLUT4 storage compartments, and converges with GLUT4 trafficking pathways during ligand-induced internalization. These results uncover a previously unrecognized connection between ACE2 and insulin-responsive vesicle dynamics, providing new insight into ACE2 function in adipocytes and its potential relevance to insulin resistance, obesity, and COVID-19 severity.

## Materials and Methods

### Materials

Dulbecco’s modified Eagle’s medium (DMEM), penicillin/streptomycin, and trypsin–EDTA were purchased from Sigma-Aldrich (St. Louis, MO, USA). Calf serum and fetal bovine serum (FBS) were obtained from BioWest (Nuaille, France). NanoBiT and HiBiT detection reagents were from Promega (Madison, WI, USA). Biotinylated recombinant SARS-CoV-2 Spike protein (Cat. #BT10549) was obtained from R&D Systems (Minneapolis, MN, USA). TAPI-1 (ADAM17 inhibitor) was purchased from commercial sources. Unless otherwise noted, all chemicals were of the purest grade available from Sigma Chemical or Wako Pure Chemical Industries (Osaka, Japan).

### Cell culture and adipocyte differentiation

3T3L1 fibroblasts were maintained in DMEM supplemented with 10% calf serum and 1% penicillin/streptomycin at 37°C in a humidified atmosphere containing 5% CO_2_. For adipocyte differentiation, cells were cultured in DMEM containing 10% FBS supplemented with 0.5 mM isobutylmethylxanthine (IBMX), 0.25 μM dexamethasone, and 1 μg/mL insulin for 4 days, followed by culture in DMEM with 1 μg/mL insulin for an additional 4 days [19]. Fully differentiated adipocytes (days 6–10 post-induction) were used for experiments. For insulin stimulation experiments, adipocytes were serum-starved for 3 h and treated with 100 nM insulin for 30 min (acute stimulation) or cultured in medium containing 100 nM insulin for 16 h (prolonged stimulation).

### Plasmid constructs

Expression vectors containing cDNA encoding ACE2 (pFN21AE2097), GLUT4 (pFN21AB0463), TRPV1 (pFN21AB9458) and SLC6A19 (pFN21AE0366) were purchased from Promega (Tokyo, Japan). For visualization experiments, these cDNAs were subcloned into pFC14A HaloTag CMV Flexi vector to fuse HaloTag tag at the C-terminus. Expression vectors containing cDNA encoding GLUT4-EGFP [19] were also used for fluorescent imaging studies. For NanoBiT assays, ACE2 and GLUT4 cDNAs were subcloned into pFC34K (LgBiT) and pFC36 (SmBiT) vectors to generate C-terminally tagged fusion proteins. Because ACE2 is a type I transmembrane protein, tags were fused to its intracellular C-terminus. GLUT4 was also C-terminally tagged to preserve its N-terminal trafficking signals. For surface quantification experiments, the HiBiT (VSGWRLFKKIS) sequence just after the signal peptide of hACE2 at position A25, its corresponding nucleotide sequence (5′-GTGAGCGGCTGGCGGCTGTTCAAGAAGATTAGCGGAGGA-3′), with the final GGAGGA serving as a linker, was introduced into pFC14A-ACE2-HaloTag vector by overlap extension PCR. The insertion was confirmed by sequencing.

### Transfection and electroporation

3T3L1 fibroblasts were transfected using Lipofectamine 3000 according to the manufacturer’s instructions. Typically, total 2 μg plasmid DNA was used per 35-mm glass-bottom dish for imaging or per well for luminescence assays. Differentiated adipocytes were transfected by electroporation under low-voltage conditions (160 V, 950 μF) as previously described. For each electroporation, 12.5–50 μg plasmid DNA was introduced. Cells were analyzed 24–48 h after transfection [20]. Cells were plated on glass-bottom dishes for imaging experiments or white optical-bottom 96-well plates for luminescence assays.

### NanoBiT luminescence assay

Protein association in living cells was assessed using the NanoBiT system. Cells expressing LgBiT- and SmBiT-tagged constructs were cultured for 24–48 h prior to measurement. Luminescence signals were recorded at room temperature using either the Kronos AB-2550 (Atto Corporation, Tokyo, Japan) or the GloMax® Discover System (Promega, Madison, WI, USA) following addition of Nano-Glo substrate according to the manufacturer’s protocol. For insulin stimulation experiments, cells were treated with 100 nM insulin for 30 min or 16 h prior to measurement. Luminescence values were normalized to single-construct controls.

### HiBiT-based surface exposure and shedding assays

Cell surface exposure of ACE2 was quantified using the HiBiT detection system. LgBiT protein and substrate were added to intact cells expressing HiBiT-ACE2, and luminescence corresponding to extracellularly exposed ACE2 was measured immediately. For shedding assays, conditioned media were collected after 24 h incubation, and luminescence signals were measured to quantify released HiBiT-ACE2. Where indicated, cells were treated with 100 nM insulin or 1 μM TAPI-1 prior to medium collection.

### Confocal fluorescence microscopy

Subcellular localization of ACE2-Halo, SLC6A19-Halo, and GLUT4-EGFP was examined using a confocal laser-scanning microscope (Olympus FV-1000) equipped with a PlanApo 60× oil-immersion objective lens (NA 1.40). Cells were incubated with Halo-TMR ligand for 30 min, fixed with 4% paraformaldehyde in PBS, and permeabilized with 0.1% Triton X-100 when required. For Spike protein internalization experiments, cells were incubated with biotinylated recombinant S-protein for 30 min at 37°C, followed by fixation and detection with Alexa Fluor 488-conjugated streptavidin. The acquired TIFF images were processed using ImageJ software (NIH, Bethesda, MD, USA). Quantitative co-localization analysis was performed with Intensity Correlation Analysis plug-in of ImageJ [21].

### Image analysis and quantification of fluorescence distribution

Fluorescence image analysis was performed using Fiji/ImageJ (NIH). All images were processed using identical parameters within each experiment. Background signal was removed using the Subtract Background function (rolling ball radius, 40 pixels). Individual cells were manually defined as regions of interest (ROIs), and fluorescence intensity measurements were obtained within each ROI across all channels.

Line scan analysis was performed by drawing a straight line across individual cells using the Line tool in Fiji/ImageJ. Fluorescence intensity profiles were obtained using the Plot Profile function and plotted as a function of distance. Identical line length and orientation criteria were applied across conditions. Colocalization analysis was performed using the Coloc 2 plugin in Fiji/ImageJ. Pearson’s correlation coefficient (R) was calculated within manually defined cellular ROIs using background-corrected images. Each data point represents one cell.

Signal heterogeneity was quantified using the coefficient of variation (CV), calculated as the standard deviation divided by the mean fluorescence intensity. Identical analysis parameters were applied to all images within each experiment. Each data point represents one cell.

### Reconstitution of insulin-responsive GLUT4 vesicle formation

To reconstitute insulin-responsive GLUT4 trafficking in fibroblasts, ACE2-Halo and GLUT4-EGFP were co-expressed with Sortilin and AS160 expression vectors [21]. Insulin-dependent redistribution and vesicular localization were analyzed by confocal microscopy as described above.

### Structural modeling

Structural models of ACE2–GLUT4 complexes were generated using AlphaFold2 (ColabFold, multimer mode) with default parameters. Model confidence was evaluated using average pLDDT scores. Protein–protein interfaces were analyzed using PDBe PISA, and membrane alignment was assessed using the OPM database. ACE2–SLC6A19 models were generated as positive controls.

### Statistical analysis

All experiments were performed at least three independent times. Data are presented as mean ± SD. Statistical analysis was performed using GraphPad Prism 8. Differences between groups were evaluated by one-way ANOVA followed by Tukey’s multiple comparison test. A p value < 0.05 was considered statistically significant. Comparisons between two groups were performed using the Mann–Whitney U test.

## Results

### Association between ACE2 and GLUT4 detected by NanoBiT luminescent assay under basal and insulin-stimulated conditions

To explore the potential physical interaction between GLUT4 and ACE2, analogous to the established ACE2–SLC6A19 complex [15], we examined their direct interaction in living cells using the NanoBiT (NanoLuc Binary Technology) system.

ACE2 is a type I transmembrane protein; therefore, SmBiT or LgBiT was fused to its intracellular C-terminus. Similarly, the C-terminus of GLUT4 was fused with either SmBiT or LgBiT, as N-terminal tagging may interfere with proper GLUT4 trafficking [22]. Among four possible tag combinations, ACE2-SmBiT and GLUT4-LgBiT produced the highest luminescent intensity compared with other combinations or single constructs (**Fig. 1A**).

**Figure 1.**
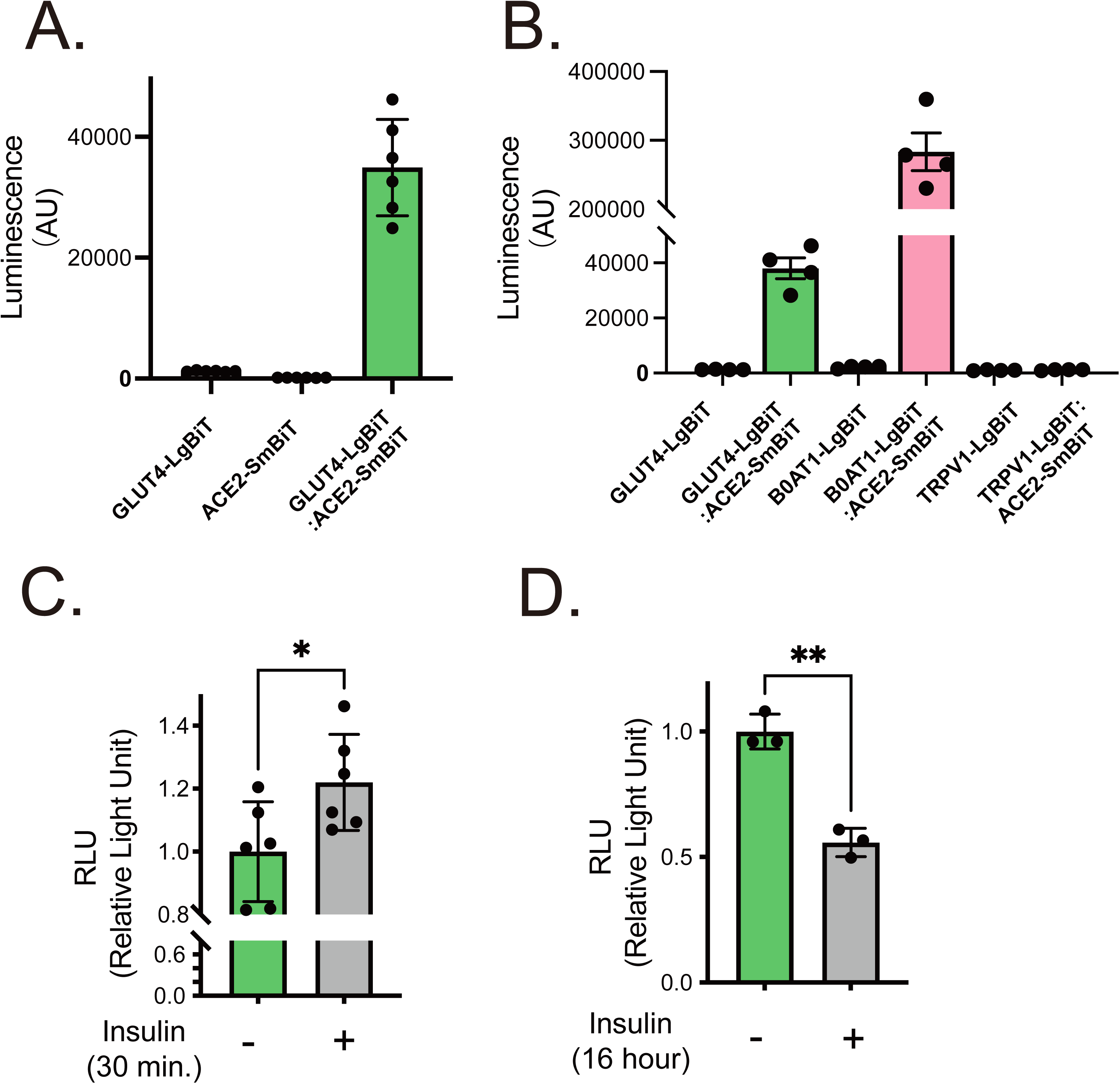
Regulated association between ACE2 and GLUT4 detected by NanoBiT assay. **(A)** Expression vectors containing cDNA encoding C-terminally LgBiT- or SmBiT-fused ACE2 (ACE2-SmBiT, ACE2-LgBiT) and GLUT4 (GLUT4-SmBiT, GLUT4-LgBiT) were transfected into 3T3L1 adipocytes and the interaction between LgBiT- and SmBiT-fused membrane proteins of interest was evaluated by NanoBiT-based luminescent intensity as described in Materials and Methods. Note that each expression vector alone displayed minimal or no luminescence. Representative data from one experiment with 4–6 wells per condition are shown. Similar results were obtained in three independent experiments. **(B)** Expression vectors for ACE2-SmBiT and either GLUT4-LgBiT, SLC6A19-LgBiT, or TRPV1-LgBiT were transfected into 3T3L1 adipocytes and the interaction between ACE2 and GLUT4 or SLC6A19 was evaluated. Minimal or no luminescence was observed with GLUT4-LgBiT, SLC6A19-LgBiT, or TRPV1-LgBiT alone. Representative data from one experiment with 4–6 wells per condition are shown. Similar results were obtained in three independent experiments. **(C)** Effect of acute insulin stimulation (100 nM, 30 min) on NanoBiT luminescence. **(D)** Effect of prolonged insulin exposure (100 nM, 16 h) on NanoBiT luminescence. For panels C and D, luminescence values were normalized to the no-insulin control after background subtraction. Data shown are from a representative experiment performed with 3–6 wells per condition. Similar results were obtained in three independent experiments. Statistical significance is indicated as follows: \**P* < 0.05, \*\**P* < 0.01.

As a positive control, ACE2–SLC6A19 interaction was evaluated and generated significantly higher luminescence. Conversely, ACE2-SmBiT and TRPV1-LgBiT, an unrelated plasma membrane protein with no known interaction with ACE2 ([23]), yielded no detectable luminescence signal (**Fig. 1B**). Notably, the luminescence intensity of the ACE2–SLC6A19 complex was approximately eight-fold higher than that of ACE2–GLUT4, indicating that although ACE2 shows proximity-based association with GLUT4 in living cells, this interaction is less robust and/or occurs at lower frequency compared with the canonical ACE2–SLC6A19 complex.

To assess insulin-dependent modulation, we examined acute (30 min) and chronic (16 h) insulin stimulation. Acute insulin slightly but significantly increased luminescence (**Fig. 1C**), whereas prolonged insulin exposure reduced it (**Fig. 1D**). These findings suggest that ACE2–GLUT4 proximity is dynamically regulated by insulin signaling.

### Subcellular localization of ACE2 and regulation of surface exposure and shedding in adipocytes

To examine the intracellular distribution of ACE2 in adipocytes, we performed confocal fluorescence imaging of differentiated 3T3L1 cells co-expressing ACE2-Halo and GLUT4-EGFP. Under basal conditions, ACE2 localized to both the plasma membrane and intracellular vesicular structures, with substantial overlap observed within GLUT4-positive intracellular compartments (**Fig. 2A**). In contrast, GLUT4 was largely undetectable at the plasma membrane under basal conditions. Insulin stimulation (100 nM, 30 min) induced redistribution of GLUT4 toward the plasma membrane (**Figs. 2A**). Line-scan analysis across representative intracellular regions revealed overlapping fluorescence peaks of ACE2 and GLUT4 under basal conditions, whereas minimal overlap was detected at the plasma membrane. Following insulin stimulation, coincident fluorescence signals appeared at the plasma membrane (**Fig. 2B**, *arrows*), consistent with GLUT4 translocation (**Fig. 2A**). Quantitative colocalization analysis using Pearson’s correlation coefficient confirmed significant overlap between ACE2 and GLUT4 under basal conditions. Insulin treatment further increased the colocalization index, reflecting redistribution of GLUT4 from intracellular compartments to the plasma membrane, where ACE2 was already present (**Fig. 2D**).

**Figure 2.**
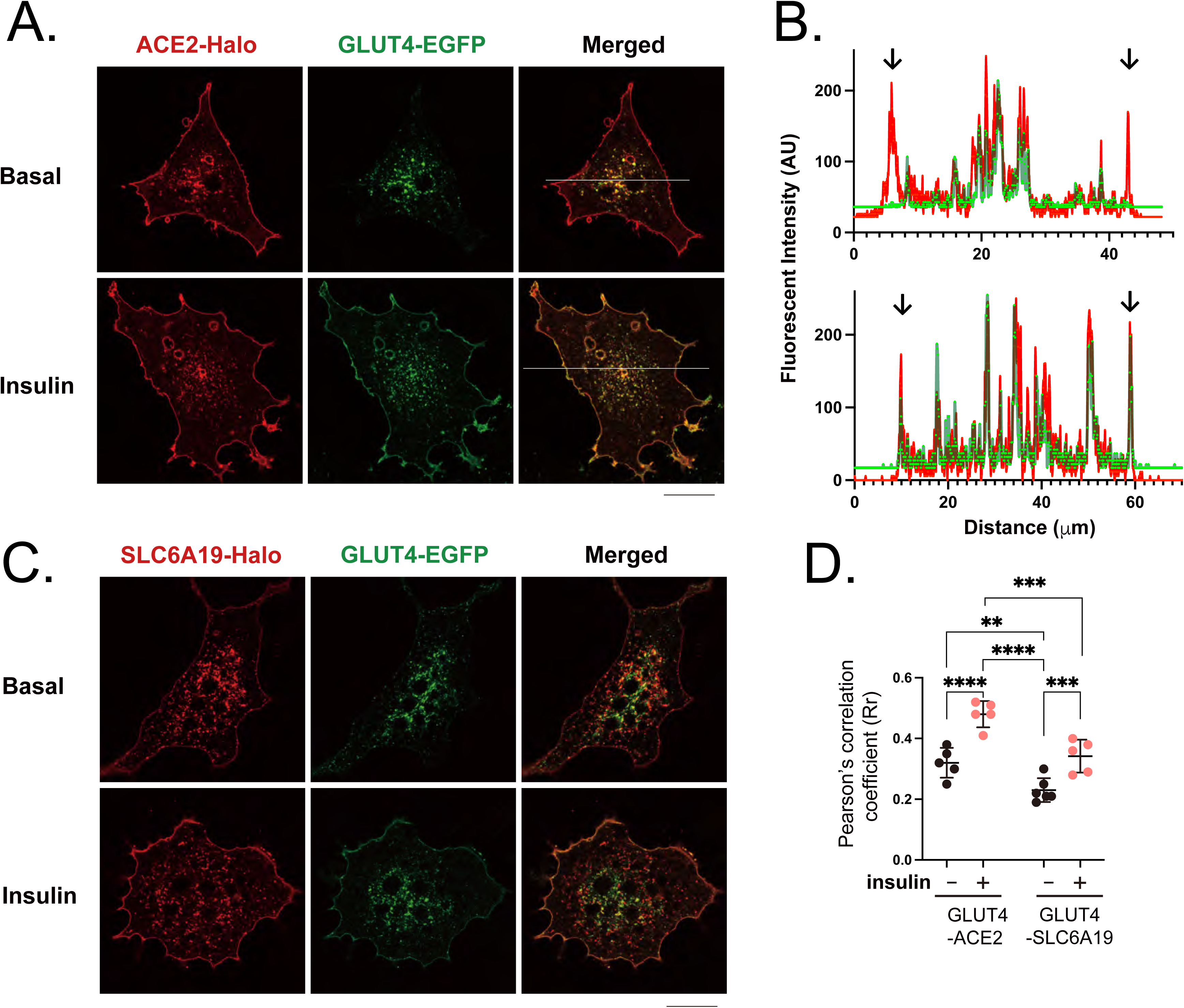
Subcellular localization and insulin-responsive regulation of ACE2 in 3T3L1 adipocytes. (A) Confocal images of differentiated 3T3L1 adipocytes co-expressing ACE2-Halo (red) and GLUT4-EGFP (green) under basal conditions or after insulin stimulation (100 nM, 30 min). Merged images are shown at right. Scale bar, 20 μm. Representative images are shown from one of three independent experiments in which more than 30 cells were analyzed per experiment. (B) Line-scan fluorescence intensity profiles corresponding to the regions indicated in (A). Red and green traces represent ACE2-Halo and GLUT4-EGFP fluorescence, respectively. Arrows indicate plasma membrane regions. Profiles were obtained from the representative cells shown in (A) and showed similar patterns across independent experiments. (C) Confocal images of adipocytes co-expressing SLC6A19-Halo (*red*) and GLUT4-EGFP (*green*) under basal or insulin-stimulated conditions. Scale bar, 20 μm. Representative images are shown from one of three independent experiments in which more than 30 cells were analyzed per experiment. (D) Quantification of colocalization using Pearson’s correlation coefficient (R) calculated from intracellular regions. Each dot represents one cell from a representative experiment. Similar results were obtained in three independent experiments. Data represent mean ± SD of analyzed cells, and statistical significance is indicated as follows: \*\**P* < 0.01, \*\*\**P* < 0.005, \*\*\*\**P* < 0.001.

To compare localization patterns with another ACE2-associated protein, SLC6A19 was ectopically introduced as an experimental reference protein, although it is not normally present in adipocytes (**Fig. 2C**). Similar to ACE2, SLC6A19-Halo was localized to both the plasma membrane and intracellular compartments under basal condition; however, its intracellular distribution showed limited intracellular colocalization with GLUT4. Insulin stimulation modestly increased apparent overlap, primarily reflecting GLUT4 redistribution to the plasma membrane rather than stable intracellular association (**Figs. 2C and 2D**). Interestingly, despite evident intracellular colocalization of ACE2 and GLUT4 by confocal microscopy, the ACE2–GLUT4 pair generated relatively lower NanoBiT luminescence compared with ACE2–SLC6A19. In contrast, SLC6A19 showed limited overlap with GLUT4 in compartments where ACE2 was present, yet produced stronger luminescence signals when co-expressed with ACE2 (**Fig. 1**). These findings indicate that luminescence intensity does not necessarily correlate with the degree of spatial colocalization detected by microscopy.

Because confocal fluorescence imaging alone did not allow clear discrimination of possible insulin-induced ACE2 translocation, we quantitatively evaluated ACE2 plasma membrane exposure using HiBiT-ACE2 containing an extracellular HiBiT tag. Surface-exposed HiBiT-ACE2 was detected by extracellular complementation with LgBiT protein in the presence of the luminescent substrate furimazine. Insulin stimulation resulted in a slight but significant increase in plasma membrane-associated ACE2, as measured by luminescence intensity (**Fig. 3A**). These findings suggest that a fraction of ACE2 present in intracellular GLUT4 storage vesicles translocates to the cell surface in response to insulin. A rim fluorescence assay using GLUT4-EGFP confirmed robust insulin-stimulated GLUT4 translocation and demonstrated that ACE2 overexpression did not alter GLUT4 trafficking dynamics in 3T3L1 adipocytes (**Fig. 3B**).

**Figure 3.**
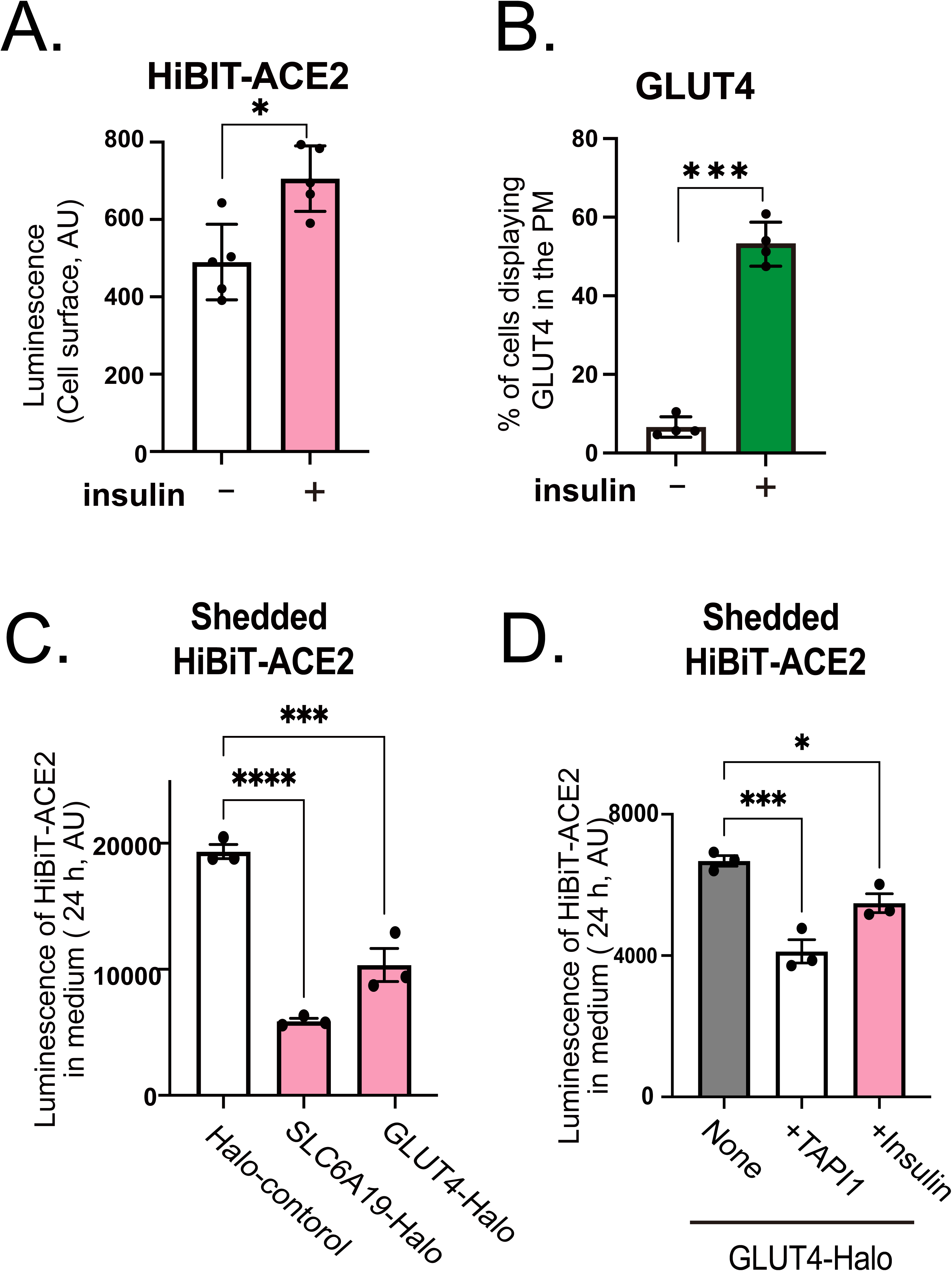
Regulation of ACE2 surface exposure and ectodomain shedding by insulin and GLUT4-associated trafficking. (A) Cell surface exposure of HiBiT-ACE2 measured by extracellular complementation with LgBiT and substrate under basal or insulin-stimulated conditions. (B) Percentage of cells displaying surface GLUT4 following insulin stimulation. (C) Quantification of shed HiBiT-ACE2 in conditioned medium (24 h) in cells co-expressing Halo control, SLC6A19-Halo, or GLUT4-Halo. (D) Effect of TAPI-1 or insulin on ACE2 shedding in GLUT4-Halo–expressing cells. Data represent mean ± SD of technical replicates from a representative experiment performed with 3–6 wells per condition. Similar results were obtained in three independent experiments. Statistical significance is indicated as follows: \**P* < 0.05, \*\*\**P* < 0.005, \*\*\*\**P* < 0.001.

We next assessed whether molecular association influences ACE2 shedding. Quantification of shed HiBiT-ACE2 in conditioned medium revealed that co-expression of GLUT4-Halo or SLC6A19-Halo significantly reduced ACE2 shedding compared with Halo control (**Fig. 3C**). As previously reported [24], treatment with the metalloprotease inhibitor TAPI-1 suppressed ACE2 shedding, indicating the involvement of a TAPI-1–sensitive protease (**Fig. 3D**). Interestingly, prolonged insulin exposure (24 h) also resulted in a significant reduction in ACE2 shedding (**Fig. 3D**). Under the same conditions, NanoBiT luminescence between ACE2 and GLUT4 was significantly decreased (**Fig. 1**), suggesting that sustained insulin stimulation alters their molecular association and/or trafficking dynamics.

Together, these findings indicate that ACE2 shedding is mediated by a TAPI-1–sensitive protease and that its shedding efficiency is modulated in the presence of exogenously expressed GLUT4 or SLC6A19. Moreover, stimuli that alter trafficking context and vesicular dynamics, such as insulin stimulation, further influence ACE2 processing in a complex manner. These observations also suggest that ACE2 shedding is determined not only by intrinsic protease activity but also by the trafficking context imposed by its molecular partners, which may in turn govern protease accessibility.

### A fraction of ACE2 is recruited into insulin-responsive GLUT4 storage vesicles upon exogenous expression of Sortilin and AS160 in fibroblasts

To further characterize subcellular localizations of GLUT4 and ACE2, we exogenously expressed both GLUT4 and ACE2 in 3T3L1 fibroblasts, which lack the mature GLUT4 trafficking system characterized by perinuclear GLUT4 retention and its insulin-responsive mobilization for GLUT4 translocation [21, 25]. Unlike fully differentiated 3T3L1 adipocytes (**Fig. 2A**), in 3T3L1 fibroblasts, ACE2-Halo and GLUT4-EGFP diffusely localized at the plasma membrane and cytoplasm, with only small and inconspicuous granular structures (**Fig. 4A**). Insulin stimulation did not noticeably induce GLUT4-EGFP translocation in 3T3L1 fibroblasts, as previously reported [21], nor did it significantly affect the localization pattern of ACE2. Pixel intensity scatter plots comparing ACE2 and GLUT4 signals revealed a compact distribution along the diagonal axis, indicating proportional signal correlation and relatively homogeneous intracellular distribution. These observations suggest that, in fibroblasts lacking the mature GLUT4 trafficking machinery [26]. ACE2 and GLUT4 remain largely diffusely distributed.

**Figure 4.**
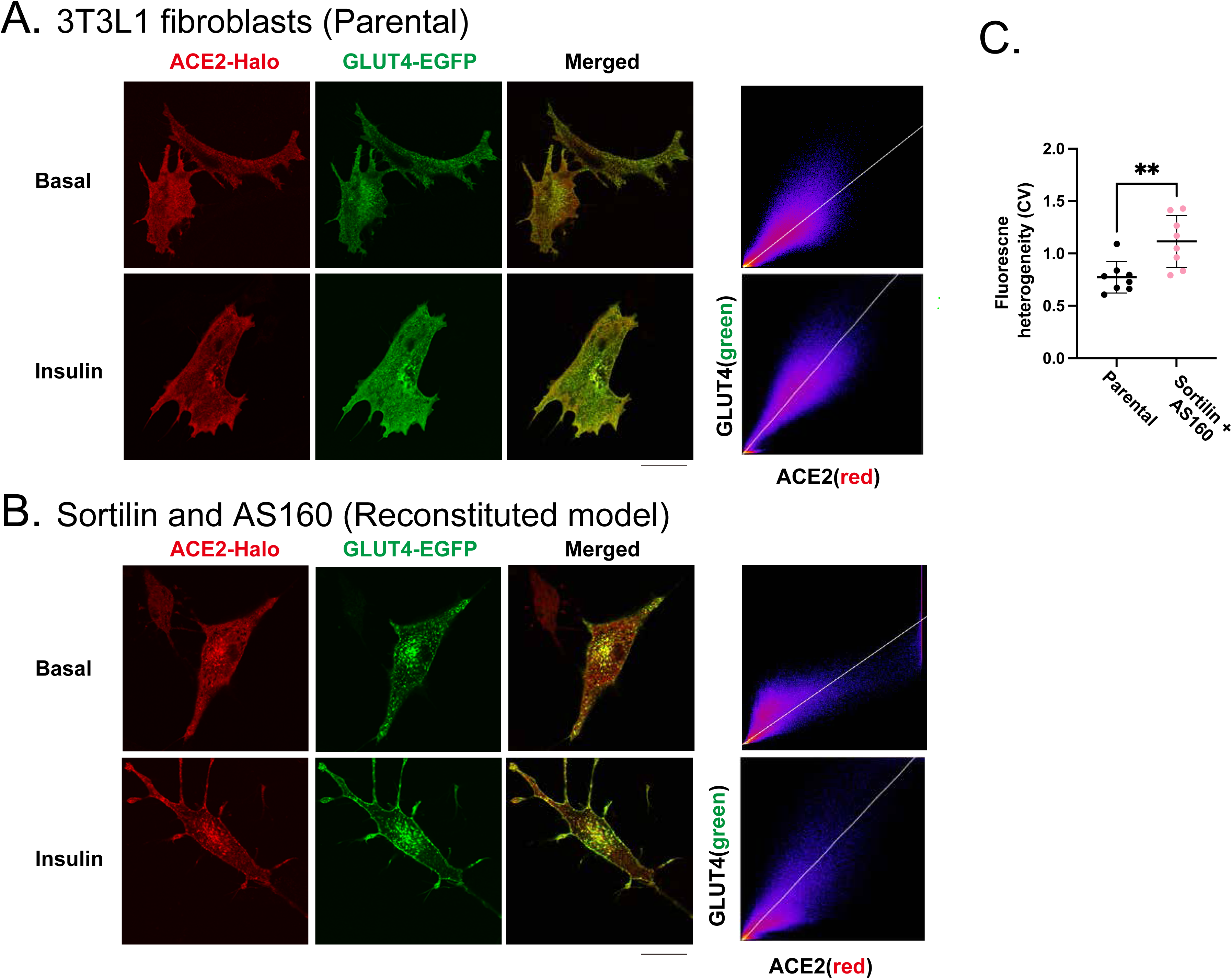
Reconstitution of insulin-responsive GLUT4 trafficking promotes vesicular co-localization of ACE2. **(A)** Confocal images of parental 3T3L1 fibroblasts expressing ACE2-Halo (*red*) and GLUT4-EGFP (*green*) under basal conditions or after insulin stimulation (100 nM, 30 min). In parental fibroblasts, both proteins exhibit largely diffuse intracellular distribution with minimal vesicular accumulation. Representative images are shown from one of three independent experiments. Scale bars, 20 μm. Corresponding pixel intensity scatter plots show compact signal correlation, indicating limited spatial clustering. **(B)** Confocal images of reconstituted insulin-responsive 3T3L1 fibroblasts co-expressing ACE2-Halo, GLUT4-EGFP, sortilin, and AS160. In these cells, GLUT4 localizes to perinuclear vesicular structures that also contain ACE2 under basal conditions and undergo insulin-responsive redistribution. Corresponding scatter plots show increased signal dispersion and high-intensity tails, consistent with heterogeneous vesicular clustering and enhanced co-distribution of ACE2 with GLUT4. Representative images are shown from one of three independent experiments. Scale bars, 20 μm. **(C)** Quantification of fluorescence heterogeneity measured as the coefficient of variation (CV = standard deviation / mean fluorescence intensity) within cellular regions of interest. Each dot represents one cell from a representative experiment. Similar results were obtained in three independent experiments. Data represent mean ± SD of analyzed cells, and statistical significance is indicated as follows: \*\**P* < 0.01.

Because partial but clear co-localization of ACE2 and GLUT4 within the perinuclear GLUT4 storage compartment was observed only in differentiated 3T3L1 adipocytes (**Fig. 2A**), but not in fibroblasts (**Fig. 4A**), we took advantage of a cell-based reconstitution model using 3T3L1 fibroblasts (Sortilin/AS160-3T3L1-cells). This model recapitulates both perinuclear segregation of GLUT4 and insulin-responsive GLUT4 redistribution through exogenous expression of Sortilin and AS160, which are both highly upregulated upon differentiation, enabling the establishment of the mature GLUT4 trafficking system [21, 25]. In Sortilin/AS160-3T3L1 cells, GLUT4-EGFP predominantly localized in vesicular structures around the perinuclear region, which also contained ACE2-Halo under basal conditions (**Fig. 4B***, upper panels*). As we previously reported, insulin stimulation induced clear translocation of GLUT4-EGFP in the AS160/Sortilin-reconstituted 3T3L1-cells (**Fig. 4B**, *lower panels*). Consistent with these observations, pixel intensity scatter plots showed a broader distribution with an extended high-intensity tail, indicative of clustered vesicular localization rather than diffuse distribution. To quantitatively assess spatial organization, fluorescence heterogeneity was measured using the coefficient of variation (CV = standard deviation / mean fluorescence intensity) within cellular regions of interest. Cells expressing Sortilin and AS160 exhibited significantly increased CV values compared with control fibroblasts, indicating enhanced spatial heterogeneity and punctate vesicular organization (**Fig. 4C**).

Taken together, while ACE2 does not impact GLUT4 localization or its insulin-responsive translocation (**Figs. 2A and 3B**), ACE2 appears to co-localize with GLUT4, and its localization at least partially relies on GLUT4 distribution. Because GLUT4 distribution changes during adipogenic differentiation (**Fig. 2A**) and can be reconstituted in fibroblasts by exogenous expression of sortilin and AS160 to restore a mature GLUT4 trafficking system (**Fig. 4B**) [21], ACE2 localization appears to vary according to GLUT4 distribution and is therefore context-dependent.

### Subcellular localization of ACE2 after S-protein treatment in 3T3L1 cells expressing ACE2-Halo

To further characterize ACE2 distribution, we examined the effect of exogenous recombinant S-protein administration on subcellular ACE2 localization in 3T3L1 fibroblasts and adipocytes (**Fig. 5**). In these experiments, biotinylated S-protein was administered and incubated for 30 min at 37 ℃, followed by detection of the cell-surface bound and endocytosed S-proteins using Alexa488-labeled streptavidin with membrane permeabilization. In the absence of S-protein, ACE2-Halo was localized diffusely at both the plasma membrane and cytoplasm, with small and inconspicuous granular structures in 3T3L1 fibroblasts (**Fig.5A***, upper panels*). However, after S-protein administration, a large number of endosomes containing both ACE2 (*red*) and endocytosed S-protein (*green*) were observed (**Fig. 5A**, *lower panels*). Signal heterogeneity was quantified using the coefficient of variation (CV), calculated as the standard deviation divided by the mean fluorescence intensity (**Fig. 5A**, *right graph*). The CV increased from 0.82 in control cells to 1.29 following S-protein treatment, indicating a marked increase in spatial heterogeneity and signal clustering.

**Figure 5.**
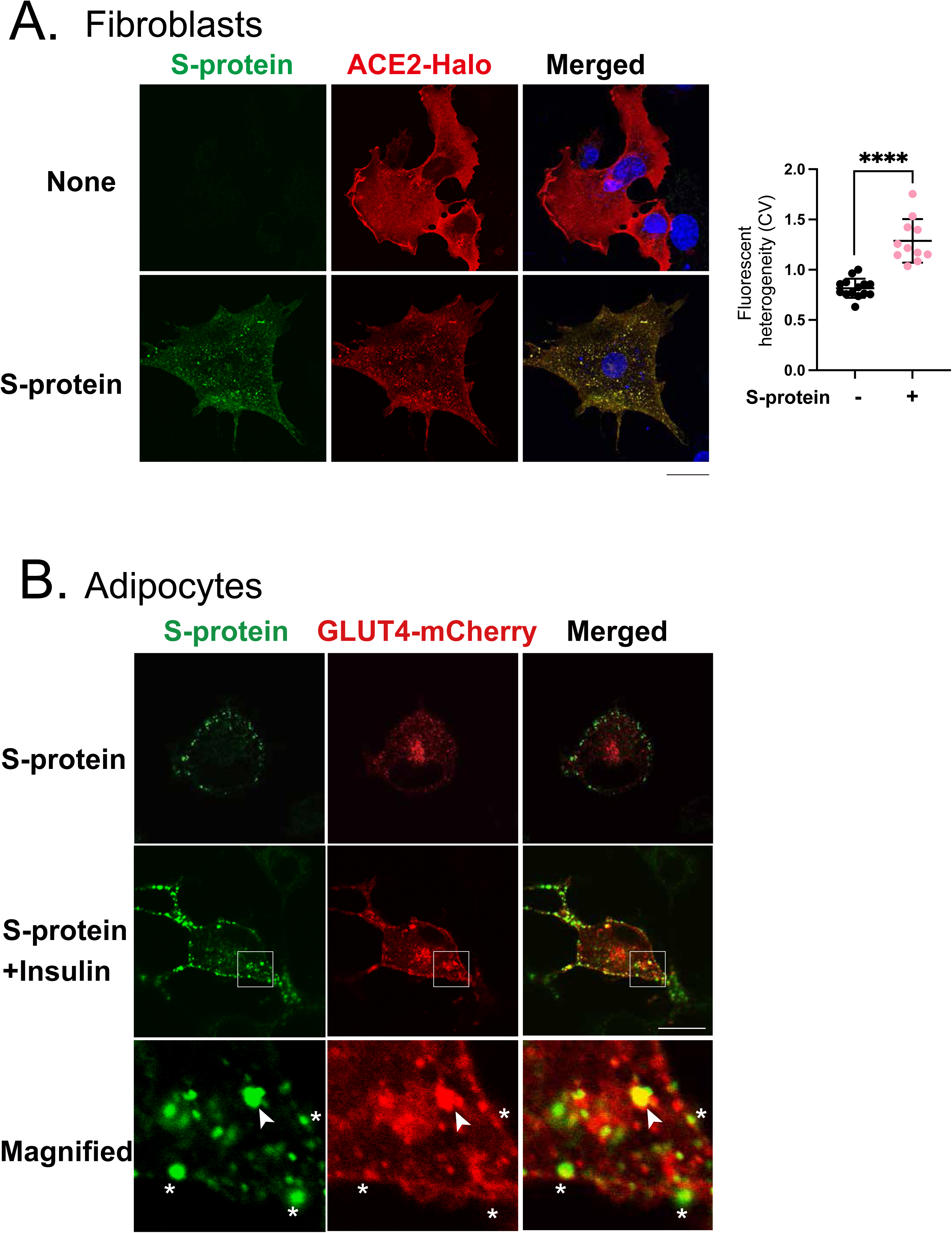
Internalization of ACE2 and trafficking relationship with GLUT4 following S-protein treatment. (A) 3T3L1 fibroblasts expressing ACE2-Halo were incubated with biotinylated recombinant S-protein (30 min, 37 °C). Internalized S-protein was detected using Alexa Fluor 488-conjugated streptavidin following permeabilization. Endosomal structures containing ACE2 and S-protein are shown. Representative images are shown from one of three independent experiments. Scale bars, 20 μm. *Right graph*: Quantification of fluorescence heterogeneity measured as the coefficient of variation (CV = standard deviation / mean fluorescence intensity) within cellular regions of interest. Each dot represents one cell from a representative experiment. Similar results were obtained in three independent experiments. Data represent mean ± SD of analyzed cells, and statistical significance is indicated as follows: \*\*\*\**P* < 0.001. (B) 3T3L1 adipocytes expressing ACE2-Halo and GLUT4-mCherry were treated with S-protein in the absence or presence of insulin (100 nM, 30 min). Internalized S-protein was detected using Alexa Fluor 488-conjugated streptavidin following permeabilization.Endosomal structures containing ACE2 and S-protein are shown. Colocalization of internalized S-protein with GLUT4-positive vesicles was assessed by confocal microscopy. *Arrowheads* indicate peripheral endosomes containing GLUT4 that also contain internalized S-protein. *Asterisks* indicate S-protein puncta located at or just beneath the plasma membrane that do not contain detectable GLUT4. Representative images are shown from one of three independent experiments. Scale bars, 20 μm.

In 3T3L1 adipocytes under basal condition, endocytosed S-protein did not reach the perinuclear GLUT4 storage compartments within the 30-min period (**Fig. 5B**, *upper panels*), suggesting that the endocytosed S-protein, presumably associated with ACE2, fails to access the perinuclear GLUT4 storage compartments. On the other hand, after co-stimulation with insulin and S-protein for 30 min, several, but not all, peripheral endosomes containing GLUT4-mCherry (*red*), which may have been exposed to the cell surface, showed colocalization with endocytosed S-protein (*green*) (**Fig. 5B**, *bottom panels, arrowheads*). Intriguingly, some puncta of S-protein at or just beneath the plasma membrane did not contain any GLUT4 proteins (**Fig. 5B**, *bottom panels, asterisks*). Taken together, these observations suggest that GLUT4 and ACE2 seem to follow distinct endocytic routes at the plasma membrane but eventually converge in early/sorting endosomes, possibly for further trafficking, which may vary based on the ligand-binding status of ACE2, including interaction with S-protein. Moreover, it is likely that GLUT4 released from its storage compartments after insulin stimulation can co-localize with ACE2 present at the plasma membrane and/or within endosomes.

### Analysis of ACE2–GLUT4 interactions using AlphaFold2 and membrane orientation modeling

To gain structural insight into the molecular interaction between ACE2 and GLUT4, we generated structural models of the ACE2–GLUT4 complex using AlphaFold2 (ColabFold) in multimer mode with default parameters. The predictions consistently yielded dimeric assemblies with high confidence (average pLDDT score >80), supporting the potential formation of an ACE2–GLUT4 complex (**Fig. 6A**). Interface analysis using PDBe PISA (Protein Interfaces, Surfaces and Assemblies) [27] identified multiple stabilizing interactions at the predicted dimer interface, including hydrogen bonds and electrostatic contacts. In particular, two salt bridges, Asp720–Lys50 and Lys619–Glu325, were reproducibly detected at the ACE2–GLUT4 interface, suggesting a defined ionic interaction network that may stabilize heterodimer formation (**Fig. 6D**). These contacts were retained in higher-order tetrameric assemblies predicted by AlphaFold2, indicating that interface geometry alone does not discriminate between oligomeric states.

**Figure 6.**
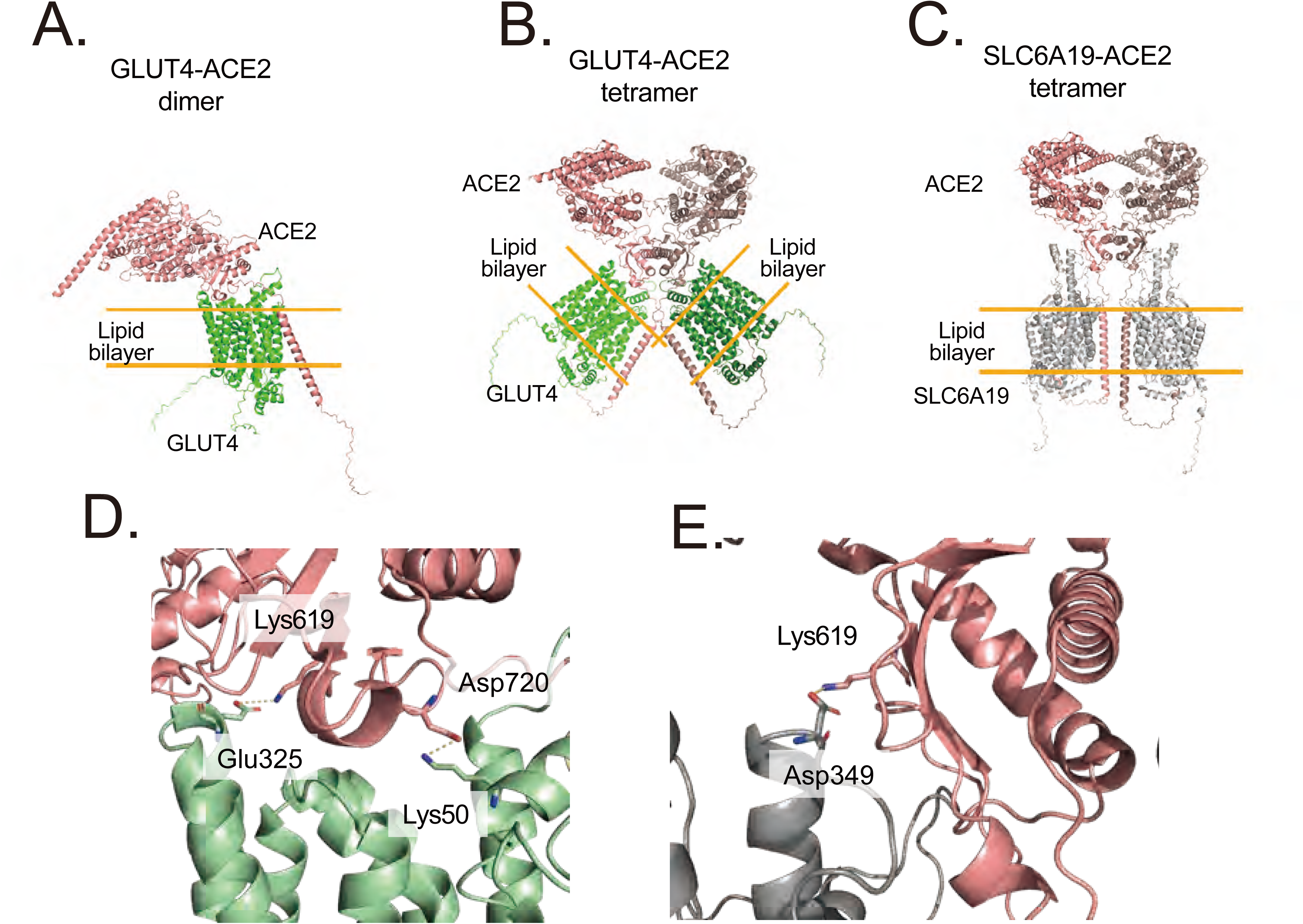
Structural analysis of ACE2 and GLUT4 interactions using AlphaFold2 and PPM membrane location prediction. (A) AlphaFold2 multimer-mode modeling generated several potential dimeric structures of the ACE2-GLUT4 complex (ACE2: pink, GLUT4: green), each with a high-confidence average pLDDT score (>80). The figure displays the structure with the highest score. (B) OPM database analysis indicated that the dimeric ACE2-GLUT4 complex aligns well with the lipid bilayer, whereas the tetrameric form showed significant misalignment, suggesting a lower likelihood of stable integration into the membrane. (C) Comparative analysis of ACE2-SLC6A19 structures was performed as a positive control. Experimentally validated tetrameric structures of ACE2-SLC6A19 heterodimers were modeled, yielding consistent structural predictions (ACE2: pink, SLC6A19: grey). The membrane positions were predicted by PPM 3.0 Web Server (https://opm.phar.umich.edu/ppm_server). (D) Two salt bridges (Asp720-Lys50 and Lys619-Glu325) located on the interface of ACE2-GLUT4 dimer. The salt bridges were detected by PDBePISA. The same salt bridges were also observed on the dimer interface in ACE2-GLUT4 tetramer. (E) A salt bridge (Lys619-Asp349) found on the dimer interface of ACE2-SLC6A19 heterodimers. The salt bridge was formed between Lys619 and Asp349 found on the extended helix of SLC6A19, which is missing in GLUT4. It is known that the crystal structure of ACE2-SLC6A19 shows a similar heterotetramer and a similar salt bridge (Lys676-Asp322) was also found on the interface, indicating this ACE2-SLC6A19 model is considered reasonable.

Because AlphaFold confidence metrics do not explicitly account for membrane constraints, we next evaluated the structural compatibility of predicted assemblies with lipid bilayers using the OPM (Orientations of Proteins in Membranes) database. Membrane orientation analysis revealed that the dimeric ACE2–GLUT4 complex exhibited favorable alignment with the lipid bilayer, whereas the tetrameric model showed pronounced misalignment relative to the membrane plane (**Fig. 6B**). This discrepancy indicates that membrane geometry imposes an additional structural constraint favoring the dimeric configuration despite comparable prediction confidence scores.

To validate this analytical framework, we applied the same modeling strategy to the ACE2–SLC6A19 complex, which has been experimentally demonstrated to form a tetramer composed of ACE2–SLC6A19 heterodimers [15]. In this control model, PDBe PISA analysis identified a distinct salt bridge (Lys619–Asp349) formed between Lys619 of ACE2 and Asp349 located within an extended helix of SLC6A19, a structural feature absent in GLUT4 (**Fig. 6E**). Consistent with experimental structures, the tetrameric ACE2–SLC6A19 assembly displayed favorable membrane alignment in OPM analysis, supporting the validity of the membrane-orientation–based evaluation.

Functionally, NanoBiT assays showed that luminescence signals from the ACE2–SLC6A19 interaction were approximately eightfold higher than those observed for ACE2–GLUT4 (**Fig. 1**), consistent with formation of a more stable higher-order assembly in the SLC6A19 complex. Taken together, these structural and functional analyses support the possibility that ACE2 preferentially adopts a membrane-compatible dimeric configuration with GLUT4, whereas tetramer formation is structurally disfavored in the absence of stabilizing architectural elements such as the extended helix present in SLC6A19.

## Discussion

In this study, we identify a previously unrecognized trafficking linkage between ACE2 and the insulin-responsive GLUT4 machinery in adipocytes. We show that ACE2 dynamically associates with GLUT4-positive compartments and that a fraction of ACE2 is recruited into insulin-responsive GLUT4 storage vesicles in a manner influenced by canonical GLUT4 trafficking components [8]. This coupling integrates metabolic signaling with ACE2 regulation, including intracellular localization (**Fig. 2**) and shedding efficacy (**Fig. 3**), and suggests that adipocyte differentiation state and insulin responsiveness critically influence ACE2 function through its subcellular compartmentalization (**Fig. 4**).

A central observation supporting this model is that ACE2 and GLUT4 exhibit a regulated proximity-based association in living cells, as detected by NanoBiT assays (**Fig. 1**). Although the ACE2–GLUT4 signal was lower than that observed for the canonical ACE2–SLC6A19 pair, luminescence was clearly above negative controls and dynamically modulated by insulin stimulation. Acute insulin modestly increased NanoBiT luminescence, whereas prolonged insulin exposure reduced it (**Fig. 1C and 1D**), suggesting that the ACE2–GLUT4 association is sensitive to insulin-dependent changes in trafficking and/or protein turnover (**Figs. 2 and 3**). Importantly, these data should not be interpreted as a simple “on/off” binding event; rather, they are consistent with a context-dependent proximity/assembly relationship that can vary with subcellular localization and cellular state.

Consistent with this view, confocal imaging revealed that in fully differentiated 3T3L1 adipocytes, ACE2 localizes not only to the plasma membrane but also to intracellular vesicular structures that substantially overlap with GLUT4-positive compartments under basal conditions (**Fig. 2**). In contrast, in 3T3L1 fibroblasts, which lack a mature GLUT4 trafficking system [25], ACE2 and GLUT4 displayed diffuse distributions with only inconspicuous puncta and no prominent perinuclear storage compartment (**Fig. 4A and 4C**). Notably, reconstitution of insulin-responsive GLUT4 vesicle formation in fibroblasts by exogenous expression of Sortilin and AS160 was sufficient to generate perinuclear GLUT4-positive vesicles [21] and to partially recruit ACE2 into these compartments (**Fig. 4B and 4C**). This reconstitution analysis provides strong support for the idea that ACE2 recruitment into perinuclear vesicular compartments is not merely a passive consequence of overexpression, but instead is consistent with its association with GLUT4 and partial coupling to the insulin-responsive GLUT4 itinerary, consistent with their proximity-based association (**Fig. 1**).

A further implication of this coordinated trafficking is that insulin can modulate ACE2 surface exposure and processing. Using extracellular HiBiT tagging, we detected a slight but significant increase in ACE2 surface exposure upon acute insulin stimulation (**Fig. 3A**), consistent with the notion that a fraction of vesicular ACE2 accompanies GLUT4 during insulin-stimulated redistribution (**Fig. 2A**). At the same time, association with GLUT4 altered ACE2 shedding: co-expression of GLUT4 reduced the release of shed ACE2 into the medium, and shedding was inhibited by the metalloprotease inhibitor TAPI-1 (**Figs. 3C** and **3D**), indicating involvement of a TAPI-1–sensitive protease, likely ADAM17 [24]. These observations suggest that ACE2 shedding efficiency is influenced not only by protease activity per se but also by the trafficking route and membrane microenvironment in which ACE2 resides, both of which can be shaped by its molecular partners including GLUT4. In this framework, recruitment of ACE2 into GLUT4-linked trafficking pathways may reduce accessibility to the shedding machinery or favor membrane domains that are less permissive for proteolytic cleavage. This model highlights how insulin signaling and metabolic states, including insulin resistance, may modulate ACE2 processing and release through trafficking pathways.

Our ligand-induced internalization experiments further support partial convergence between ACE2 and GLUT4 trafficking routes. Biotinylated S-protein triggered robust ACE2-associated endocytosis in fibroblasts, generating numerous ACE2- and S-protein–positive endosomes and increasing the spatial heterogeneity of ACE2-associated signals, indicative of ligand-induced clustering and endosomal accumulation (**Fig. 5A**). In adipocytes, endocytosed S-protein did not access the perinuclear GLUT4 storage compartments for at least 30 min under basal conditions. In contrast, upon co-stimulation with insulin and S-protein, subsets of peripheral endosomes exhibited colocalization between endocytosed S-protein and GLUT4 (**Fig. 5B**). At the same time, some S-protein puncta lacked detectable GLUT4, suggesting that ACE2 and GLUT4 can enter distinct endocytic routes at the plasma membrane but subsequently converge in early/sorting endosomal compartments. Given that ACE2 endocytosis can depend on clathrin- and cholesterol-sensitive mechanisms and may vary with ligand binding [28, 29], our findings raise the possibility that the presence and trafficking state of GLUT4 further modulate ACE2 itineraries in adipocytes. Future studies will be essential to define the molecular determinants that specify when ACE2 and GLUT4 co-internalize, when they remain segregated, and how viral ligands reshape these pathways [30].

The structural modeling provides an additional mechanistic layer that helps reconcile physical association with trafficking behavior. AlphaFold2 multimer predictions supported formation of ACE2–GLUT4 dimeric assemblies with high confidence, and PDBe PISA analysis identified stabilizing ionic contacts, including salt bridges involving Lys619 of ACE2 at the predicted interface (**Fig. 6**). However, because AlphaFold confidence metrics do not explicitly incorporate membrane constraints, we further evaluated membrane compatibility using OPM analysis. This analysis suggested that the ACE2–GLUT4 dimer aligns favorably with the lipid bilayer, whereas a predicted tetrameric arrangement is substantially misaligned and thus less likely to be membrane compatible (**Fig. 6B**). Although such configurations might be accommodated in highly curved membrane environments, such as caveolae, membrane invaginations [31], or small vesicles, the markedly lower NanoBiT luminescence observed for ACE2–GLUT4 compared with ACE2–SLC6A19 (∼eight-fold difference) suggests that stable tetramer formation is unlikely to predominate under physiological conditions. Applying the same framework to the experimentally validated ACE2–SLC6A19 complex [15] supported tetrameric assembly with favorable membrane alignment and revealed an interface salt bridge between Lys619 of ACE2 and Asp349 in an extended helix of SLC6A19 that is absent in GLUT4 (**Fig. 6E**). Together, these findings suggest that ACE2 may use a conserved interfacial hotspot (including Lys619) to engage distinct transporters, while partner-specific architectural features determine whether such interactions can support higher-order assemblies compatible with the membrane environment. This provides a plausible explanation for why ACE2–GLUT4 association is detectable yet produces lower NanoBiT luminescence than ACE2–SLC6A19.

The physiological and pathological implications of these findings may extend beyond intracellular trafficking per se. ACE2 expression in adipose tissue and circulating soluble ACE2 (sACE2) levels show complex relationships with obesity and metabolic status [5, 32]. In this context, our data suggest that insulin-regulated trafficking and molecular partnership with GLUT4 can influence ACE2 surface exposure and shedding, potentially affecting the balance between membrane-bound ACE2 and sACE2. Although the contribution of adipose tissue to circulating sACE2 remains uncertain, the observation that trafficking context can modulate ACE2 shedding provides a conceptual route by which metabolic state might influence ACE2 processing and possibly susceptibility to, or severity of, SARS-CoV-2 infection. Moreover, this trafficking-based view may help interpret the context-dependent metabolic phenotypes reported in ACE2-deficient models [9, 10], where insulin sensitivity, adiposity, and glucose homeostasis vary with age, diet, and experimental conditions.

Several limitations should be acknowledged. First, our analyses rely on overexpression and C-terminal tagging (NanoBiT/Halo/HiBiT), which may influence protein proximity or trafficking behavior. Accordingly, the ACE2–GLUT4 association described here primarily reflects proximity detected using tagged constructs rather than direct demonstration of an endogenous complex. Nevertheless, the convergence of proximity assays, vesicle reconstitution experiments, and trafficking analyses consistently supports the conclusion that ACE2 can access GLUT4-associated trafficking pathways. Second, while confocal imaging and luminescence assays indicate partial recruitment of ACE2 into GLUT4-associated vesicular compartments, these approaches do not establish that ACE2 is restricted to canonical GLUT4 storage vesicles. Rather, ACE2 and GLUT4 likely share overlapping trafficking routes without necessarily co-residing in identical vesicles at all times. Finally, all experiments were conducted in 3T3L1 cells, and validation in primary adipocytes or in vivo models will be important to establish the physiological relevance of GLUT4-linked ACE2 trafficking.

In conclusion, our study identifies a previously unrecognized connection between ACE2 and insulin-responsive GLUT4 vesicle dynamics in adipocytes. ACE2 can associate with GLUT4 and access GLUT4-linked trafficking pathways in a manner influenced by insulin-responsive GLUT4 vesicle machinery. Rather than forming a constitutive complex, the ACE2–GLUT4 relationship appears to be dynamically regulated by cellular context and insulin signaling through mechanisms that govern GLUT4 trafficking, thereby influencing ACE2 localization, membrane exposure, and shedding. These findings suggest insulin-regulated GLUT4 vesicle trafficking as a potential link between metabolic state and ACE2 intracellular behavior in adipocytes. More broadly, this work provides a conceptual framework for understanding how modest and context-dependent protein interactions within specialized trafficking systems can fine-tune membrane protein fate, with implications for metabolic disorders and ACE2-associated pathophysiology, including obesity [1, 33], insulin resistance [34], and viral infection.

## Acknowledgements

The authors thank Natsumi Emoto for technical assistance. This study was supported by a grant from the Japan Society for the Promotion of Science (JSPS) under the Bilateral Joint Research Program (The Royal Society, UK) (no. SPJSBP 120235702 to MK). This study was also partially supported by a grant from the JSPS (no. 24K02874 to MK).

## Notes

### Competing Interest Statement

The authors have declared no competing interest.

